# Indigenous cattle of Sri Lanka: Genetic and phylogeographic relationship with Zebu of Indus valley and South Indian origin

**DOI:** 10.1101/2023.02.23.529662

**Authors:** LGS Lokugalappatti, Saumya Wickramasinghe, P.A.B.D Alexander, Kamran Abbas, Tanveer Hussain, Saravanan Ramasamy, Vandana Manomohan, Arnaud Stephane R. Tapsoba, Rudolf Pichler, Masroor E. Babar, Kathiravan Periasamy

## Abstract

The present study reports the population structure, genetic admixture and phylogeography of island cattle breeds of Sri Lanka viz. Batu Harak, Thawalam and White cattle. Moderately high level of genetic diversity was observed. Estimates of inbreeding for Thawalam and White cattle breeds were relatively high with 6.1% and 7.2% respectively. Genetic differentiation of Sri Lankan Zebu (Batu Harak and White cattle) was lowest with Red Sindhi among Indus valley zebu while it was lowest with Hallikar among the South Indian cattle. Global F statistics showed 6.5% differences among all the investigated zebu cattle breeds and 1.9% differences among Sri Lankan zebu breeds. The Sri Lankan zebu cattle breeds showed strong genetic relationships with Hallikar cattle, an ancient breed considered to be ancestor for most Mysore type draught cattle breeds of South India. Genetic admixture analysis revealed high levels of breed purity in Lanka White cattle with >97% zebu ancestry while significant taurine admixture was observed in Batu Harak and Thawalam cattle. Two major zebu haplogroups, I1 and I2 were observed in Sri Lankan zebu with the former predominating the later in all the three breeds. A total of 112 haplotypes were observed in the studied breeds, of which 50 haplotypes were found in Sri Lankan zebu cattle. Mismatch analysis revealed unimodal distribution in all the three breeds indicating population expansion. The sum of squared deviations (SSD) and raggedness index were non-significant in both the lineages of all the three breeds except for I1 lineage of Thawalam cattle (P<0.01) and I2 lineage of Batu Harak cattle (P<0.05). The results of neutrality tests revealed negative Tajima’s D values for both the lineages of Batu Harak (P>0.05) and White cattle (P>0.05) indicating an excess of low frequency polymorphisms and demographic expansion. Genetic dilution of native zebu cattle germplasm is a cause for concern and it is imperative that national breeding organizations consider establishing conservation units for the three native cattle breeds to maintain breed purity and initiate genetic improvement programs.

## 1. Introduction

Cattle represent a major proportion of livestock population in Sri Lanka. Cattle genetic resources in Sri Lanka comprise indigenous zebu (*Bos indicus*), exotic zebu, commercial taurine (*Bos taurus*) and various crosses of these three types. Historically, cattle rearing in Sri Lanka dates back to 543 BC with the arrival of “Indo Aryans” and since then, periodic introduction of various cattle breeds has taken place through time (Jalatge, 1986; Chandrasiri, 2004; Silva et al., 2008; Perera and Jayasuriya 2008). The ancient “archaic cattle” believed to be introduced by the early settlers are considered as probable ancestors of the locally adapted indigenous cattle (Felius, 1995). Since 1930s, a number of exotic breeds including Indo-Pakistan Zebu (Red Sindhi, Tharparkar, Sahiwal, Khillari, etc.), European/American taurine cattle (Friesian, Ayrshire, Shorthorn, Jersey etc.) and commercial composites (Australian Friesian Sahiwal (AFS), Australian Milking Zebu (AMZ), Girolando, New Zealand Kiwi etc.) have been introduced to improve the productivity of local cattle (Jalatge, 1986; Chandrasiri, 2004; Perera and Jayasuriya 2008). Apart from introgression of exotic germplasm, genetic and demographic processes over time have also shaped the genetic composition, population structure and diversity of extant cattle breeds in the country (Tilakaratne et al., 1974; Buvanendran and Mahadevan 1975; Jalatge, 1986; Chandrasiri, 2004).

The indigenous zebu represents more than 50% of the total cattle population, counting 1.08 million heads in the country (FAOSTAT, 2019). Although milk productivity per animal is low (2-4 liters per day), the indigenous zebu is known for their characteristics such as draught power, ability to subsist on poor quality feed, adaptability to tropical heat and resistance to diseases (Chandrasiri, 2004; Silva et al., 2008). At least four distinct breeds/types of Sri Lankan indigenous zebu have been described (Chandrasiri, 2004; Silva et al., 2010; Shanjayan and Lokugalappatti 2015). “Batu Haraka” or the “Lanka cattle” has varying shades of black to brown coat color and are distributed in the country’s dry zone. They do not possess a prominent hump or dewlap and are small and compact in body size with an average adult weight of 160 kg (Silva et al., 2008). “White cattle” or “Thamankaduwa” is predominantly found in the Eastern province and has white to grey coat color (Shanjayan and Lokugalappatti 2015). “Thawalam cattle” is an isolated population of indigenous zebu which are used as pack animals in the Central and Uva provinces (Chandrasiri, 2004). “Cape” or “Hatton” was a synthetic breed evolved by crossing taurine males (brought from Cape of Good Hope in South Africa) with local female zebu cattle. This breed was a small-sized locally adapted dairy type breed but got deteriorated subsequently due to lack of scientific breeding programs and is now believed to be extinct (FAO, 2000; Chandrasiri, 2004; National Livestock Breeding Policy guidelines and strategies for Sri Lanka, 2010). Cross breeding zebu with exotic breeds have resulted in genetic dilution of indigenous Sri Lankan cattle (Chandrasiri, 2004). As a result, the zebu breeds are now threatened/endangered due to significant genetic erosion and loss of potentially significant gene combinations and variability.

Although details of morphological and phenotypic characteristics, production system environment and management practices of indigenous Sri Lankan cattle is available, information on their diversity, genetic distance, admixture and phylogeography is limited (Silva et al., 2010). Evaluation of diversity within and between breeds provides insights into population structure and genetic relationships necessary for establishing priorities and strategies for conservation, genetic improvement and sustainable utilization. Molecular tools such as autosomal short tandem repeat DNA markers have been helpful for estimations of diversity, genetic differentiation and admixture (Canon et al., 2006; Vahidi et al., 2014; Ajmone-Marsan et al., 2014). Extra-nuclear mitochondrial DNA D-Loop polymorphisms (maternally inherited) are used as the markers of choice for domestication studies, including assessment of maternal lineages and their geographic origins (Joshi et al., 2004; Naderi et al., 2007; Liu et al., 2009). Hence, the present study was undertaken with the objectives of (i) estimating genetic diversity and relationship of indigenous Sri Lankan cattle with zebu of Indus valley and South Indian origin (ii) evaluating population structure and genetic admixture of Sri Lankan cattle (iii) identifying mitochondrial DNA maternal lineage and assess phylogeography of Sri Lankan native cattle.

## 2. Material and methods

### 2.1. Animal ethics statement

Blood samples were collected from jugular vein in EDTA vacutainer tubes. Sampling was performed by local veterinarians in the respective native breed tracts following the standard good animal practice. The blood samples were collected as part of routine veterinary surveillance and consent was obtained from farmers for the usage of samples in the study. Therefore, no further license from the “Institutional Committee for Care and Use of Experimental Animals” of the University of Peradeniya, Sri Lanka, Virtual University of Pakistan and the Joint FAO/IAEA Division, International Atomic Energy Agency, Vienna, Austria was required.

### 2.2. Sampling and genomic DNA extraction

Blood samples were collected from 349 cattle representing six Sri Lankan and three Indus valley zebu breeds. The number of samples collected from each of the breeds were: 51 from Batu Harak, 51 from White cattle, 25 from Thawalam, 53 from Holstein-Friesian, 27 from Ayrshire, 33 from Jersey, 11 from Frisian-Jersey crosses, 30 from Red Sindhi, 38 from Sahiwal and 30 from Tharparkar. Samples from Red Sindhi, Tharparkar and Sahiwal were collected from Indus Valley regions located in Pakistan. Additionally, short tandem repeat genotype and sequence data on two South Indian breeds, Hallikar (36) and Kangayam (51) generated in a previous study (Manomohan et al., 2021) was included for comparative analysis. A map indicating the sampling locations of Sri Lankan zebu cattle is provided in Figure 1. Briefly, a stratified random sampling procedure was followed to collect samples from unrelated cattle, based on the information provided by local farmers. DNA was extracted from whole blood using MasterPure DNA Purification Kit (Biozym, Illumina Inc, USA). DNA quality and quantity estimation was done by agarose gel electrophoresis and spectrophotometry. DNA samples were then stored at -20°C until further processing.

**Figure 1.**
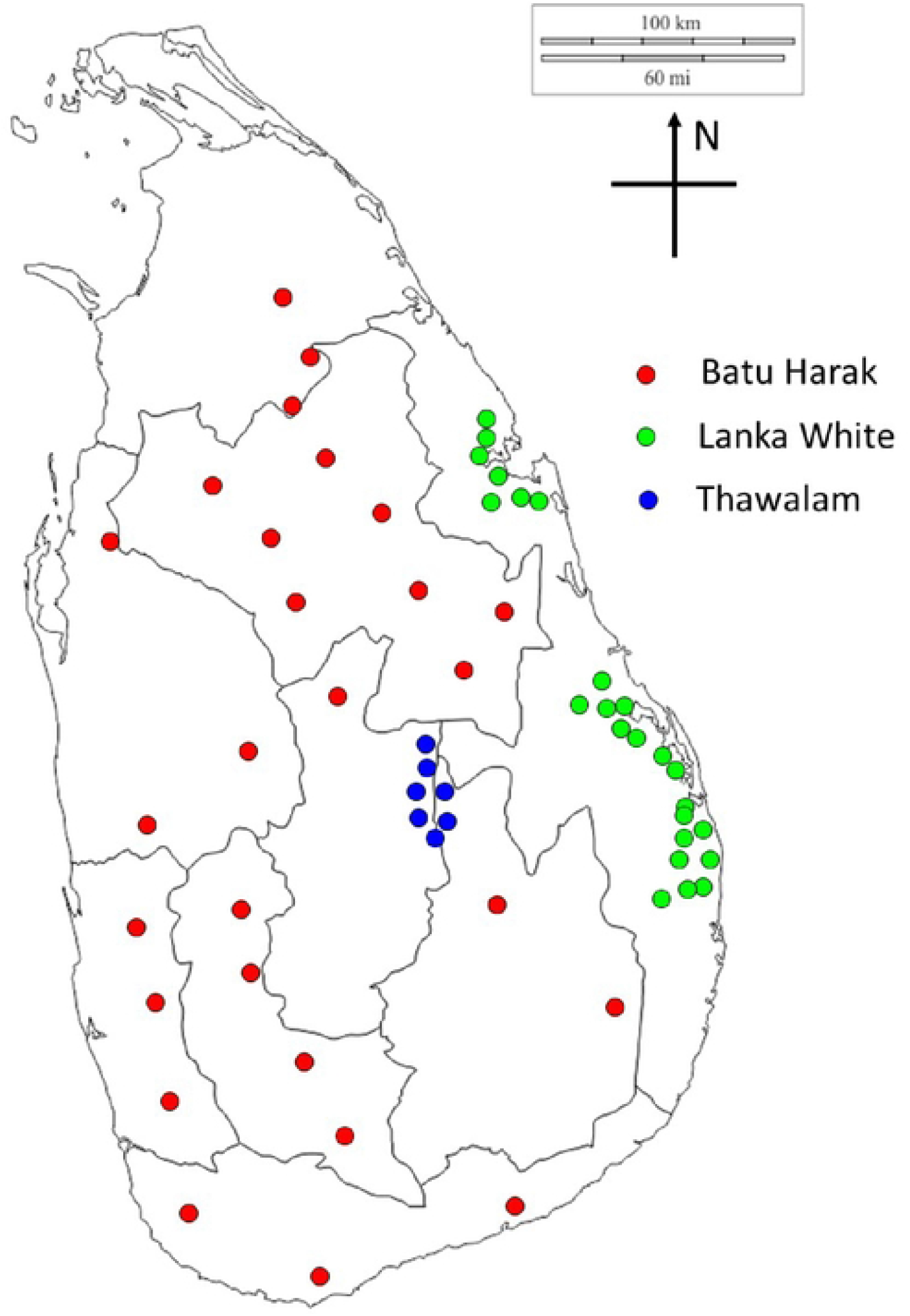
Geographical distribution of Sri Lankan zebu cattle breeds included in the study

### 2.3. Microsatellite genotyping and mitochondrial DNA sequencing

27 FAO recommended microsatellite markers (FAO, 2011) with forward primers conjugated to one of the three fluorescent dyes (FAM, HEX and ATTO550) were used for genotyping. The details of microsatellite markers, dye conjugated, annealing temperature, allele size range and PCR conditions are described elsewhere (Manomohan et al., 2021). The PCR products were capillary electrophoresed after multiplexing in an automated DNA analyzer ABI3100 (Applied Biosystems, USA) with ROX500 (Applied Biosystems, USA) as an internal lane control. The allele size data for each sample was extracted using GeneMapper v.4.1 software (Applied Biosystems, USA). The mitochondrial D-Loop region was amplified using the primers BTMTD1-F (5’ AGGACAACCAGTCGAACACC 3’) and BTMTD1-R 5’ (GTGCCTTGCTTTGGGTTAAG 3’) reported by Manomohan et al., (2021). The amplicon included complete D-loop (control region) flanked by partial cytochrome B, complete t-RNA-Thr and complete t-RNA-Pro coding sequences at 5’ end while partial t-RNA-Phe sequence flanked the 3’end. Polymerase chain reaction was performed in a total reaction volume of 40µl with the following cycling conditions: initial denaturation at 95°C for 15 min followed by 30 cycles of 95°C for 1 min; 58°C for 1 min; 72°C for 1 min with final extension at 72°C for 10 min. Purified PCR products were sequenced using Big Dye Terminator Cycle Sequencing Kit (Applied Biosystems, U.S.A) on an automated Genetic Analyzer ABI 3100 (Applied Biosystems, U.S.A).

### 2.4. Statistical analysis of data

To perform data quality control, the microsatellite genotypes were pruned for (i) missingness (ii) null alleles and (iii) deviations from selective neutrality. Individuals with >20% missing data (missing five or more genotypes out of 27 investigated loci) were excluded from the analysis. The presence of null alleles in the dataset was checked using MicroChecker version 2.2.3 (Oosterhout et al., 2004). The neutrality of the microsatellites used in this study was evaluated by comparing the markers against neutral expectations in a distribution of F_ST_ vs. heterozygosities under an island model of migration using LOSITAN version 1 (Antao et al., 2008). Basic diversity indices including observed number of alleles, observed and expected heterozygosity and pairwise Nei’s genetic distances were calculated using Microsatellite Analyzer (MSA) version 3.15 (Dieringer and Schlötterer, 2003). Pairwise Nei’s genetic distance was utilized to construct the NeighborNet tree using SplitsTree version 4.14.6 (Huson and Bryant, 2006). F statistics for each locus (Weir and Cockerman, 1984) was calculated and tested using the program FSTAT version 2.9.3.2. The exact tests for Hardy-Weinberg Equilibrium to evaluate both heterozygosity deficit and excess at each marker locus in each population (HWE) were performed using GENEPOP v4.0.9 software (Rousset 2008). Analysis of molecular variance (AMOVA) was performed using ARLEQUIN version 3.1 (Excoffier et al., 2005). Non-metric multidimensional scaling display of pairwise F_ST_ was conducted using SPSS version 13.0 (SPSS Inc, Chicago, IL, USA). The extent of population sub-structuring was further explored using STRUCTURE (Pritchard et al., 2000) with the assumption of different clusters K = 2 to K = 8. The number of burn in periods and MCMC repeats used for these runs was 200000 (Pritchard et al., 2000). The optimal ‘K’ was identified based on ΔK, the second order rate of change in LnP(D) following the procedure of Evanno et al., (2005). Mitochondrial DNA sequences were checked and edited by using Codon Code Aligner version 3.7.1. (CodonCode Corporation, Centerville, MA, USA). A total of 212 sequences generated in the present study was deposited to NCBI-GenBank (accession nos. MZ501954-MZ502165). Additionally, 87 mtDNA sequences from two South Indian cattle breeds, Hallikar (NCBI-GenBank accessions MW319954-MW319988; MW320341) and Kangayam (NCBI-GenBank accessions MW319989-MW320038; MW320342) were included for comparative analysis (Manomohan et al., 2021). Mitochondrial DNA diversity parameters including number of polymorphic, singleton and parsimony informative sites, nucleotide diversity, haplotype diversity and average number of nucleotide differences were calculated using DnaSP, version 4.10 (Rozas, 2009). Haplotype frequency, haplotype sharing and pairwise F_ST_ among studied cattle populations were calculated using ARLEQUIN 3.5.3 (Excoffier and Lischer, 2010). Pairwise F_ST_ derived from mtDNA haplotype frequency were utilized to perform non-metric multidimensional scaling (MDS) analysis using SPSS version 13.0. MEGA version 6.0 (Tamura et al., 2013) was utilized to identify optimal DNA substitution model and construct maximum-likelihood phylogeny. Reference sequences from various taurine (T1-T5) and indicine (I1 and I2) maternal lineages as mentioned in Manomohan et al., (2021) were used to construct phylogeny. HKY+I+G was identified as the optimal model for the current sequence dataset based on Akaike information criterion (AIC). To examine the possibility of a past demographic expansion, a mismatch analysis was performed by calculating frequency distributions of pairwise differences between sequences using ARLEQUIN 3.1. The tests for selective neutrality viz. Tajima’s D and Fu’s F_S_ were conducted using ARLEQUIN 3.1. Median Joining (MJ) network of Sri Lankan cattle haplotypes was reconstructed using NETWORK 4.5.1.2 (Bandelt et al., 1999).

### 3. Results

#### 3.1. Genetic variability in Sri Lankan native zebu cattle

The basic diversity measures estimated in Sri Lankan native cattle breeds are presented in Table 1. The mean number of observed alleles was high in Sri Lankan native zebu breeds and ranged from 6.96 (Thawalam) to 8.26 (Batu Harak). The mean observed heterozygosity was 0.713, 0.668 and 0.665 in Batu Harak, Sri Lankan White and Thawalam cattle respectively while the expected heterozygosity was 0.749, 0.720 and 0.708 respectively. The overall mean inbreeding estimate (F_IS_) among all the investigated cattle breeds was 0.054±0.016. Among zebu cattle alone, the overall F_IS_ was 0.063±0.020 and ranged from -0.053±0.023 (TGLA126) to 0.415±0.053 (HEL5) across different loci (Supplementary Table ST1). The mean overall F_IS_ was higher than the values reported for East Indian (Sharma et al., 2013) and Pakistani (Hussain et al., 2016) cattle breeds while lower than reported for northwestern Indian (Sodhi et al., 2011), Turkish (Özşensoy and Kurar, 2014) and Niger (Grema et al., 2017) cattle breeds. In general, F_IS_ estimates were positive at 16 of the investigated loci indicating heterozygosity deficit while remaining six loci showed evidence for heterozygosity excess (Supplementary Table ST1). The breed wise mean inbreeding estimate was 0.048, 0.072 and 0.061 in Batu Harak, White cattle and Thawalam respectively.

To further test the level of heterozygosity deficit, a total of 324 breed x locus combinations were tested for Hardy Weinberg equilibrium, of which 62 (19.14%) deviated significantly (P<0.05). Among the Sri Lankan native cattle 17 breed x loci combinations (20.99%) deviated significantly (P<0.05) due to heterozygosity deficit. The percent HWE deviation observed in the study is comparable to Brazilian (19.5% breed x locus combinations; Egito et al., 2007) and Indian zebu cattle (17.3% breed x locus combinations; Sharma et al., 2015). This could be due to factors such as presence of null alleles, selective forces operating at certain loci, intense artificial selection and/or use of few breeding bulls in the region, assortative mating, population sub-division and inbreeding within cattle herds. The MicroChecker analysis of Sri Lankan cattle genotypes at all loci revealed no significant presence of null alleles. Further, to evaluate the influence of selection on the microsatellite loci under study, a F_ST_ outlier approach was followed. Loci with an unusually high F_ST_ are putatively under directional selection, while loci with low F_ST_ value are considered to be under stabilizing selection. Among the 27 loci investigated, ETH152 was deviating from selective neutrality due to positive selection while TGLA53 was deviating significantly possibly under the influence of balancing selection (Supplementary Figure SF1). Both these loci deviated from selective neutrality when assumed under both infinite allele and step wise model of mutations and have been reported to be candidates influenced by selection (Smith et al., 2001; Li et al., 2010). Hence, ETH152 and TGLA53 were removed from further analysis in the study.

#### 3.2. Genetic differentiation among cattle breeds

Global F-statistics was performed to estimate the genetic differentiation among Sri Lankan zebu, Indus valley zebu, South Indian zebu and commercial taurine breeds, the results of which are presented in Supplementary Table ST1. The global F_ST_ was estimated at 0.145±0.010 indicating 14.5% of the total variation being explained by between breed differences and 85.5% of the variation due to within breed differences. The between breed differences was estimated at 6.5% when all the zebu breeds were considered while it was only 1.9% when Sri Lankan zebu (Batu Harak, White cattle and Thawalam) alone were considered. The pairwise F_ST_ estimates among zebu cattle breeds ranged from 0.015 (LWC-LBH; LWC-LTM) to 0.085 (LTM-IKA) (Table 2). Genetic differentiation of Sri Lankan Zebu (Batu Harak and White cattle) was lowest with Red Sindhi among Indus valley breeds while it was lowest with Hallikar among the South Indian breeds. Thawalam cattle was showing relatively higher pairwise F_ST_ estimates with both Indus valley and South Indian breeds. The pairwise Nei’s genetic distance estimates showed a similar trend among Sri Lankan, Indus valley and South Indian zebu cattle (Table 2). The NeighborNet tree derived from pairwise Nei’s genetic distance revealed distinct clustering of commercial taurine cattle and zebu breeds (Sri Lankan, Indus valley and South Indian zebu) as expected. Within the taurine cluster two distinct sub clusters could be recognized one with Ayrshire and Holstein-Friesian and the other with Friesian X Jersey crossbreds and purebred Jersey. This clearly positioned the Friesian x Jersey crossbreds between purebred Jersey and Holstein-Friesians. Among the zebu, Sri Lankan White and Thawalam cattle clustered with Hallikar cattle (South Indian zebu) while Batu Harak cattle clustered with Indus valley (Sahiwal) breeds (Figure 2). Genetic relationship among investigated cattle breeds was elucidated by implementing multidimensional scaling display of pairwise F_ST_. The normalized raw stress was 0.0019 indicating excellent goodness of fit and representation of data in the model Kruskal et al., (1964). The Tucker’s coefficient of congruence was estimated to be 0.999 implying >99% of variance in the model being accounted by the two predicted dimensions (Moroke 2014). The MDS plot showed distinct clusters of zebu and commercial taurine cattle with all the three native Sri Lankan cattle (White cattle, Batu Harak and Thawalam) placed in between zebu breeds of Indus valley and South Indian origin (Figure 3).

**Figure 2.**
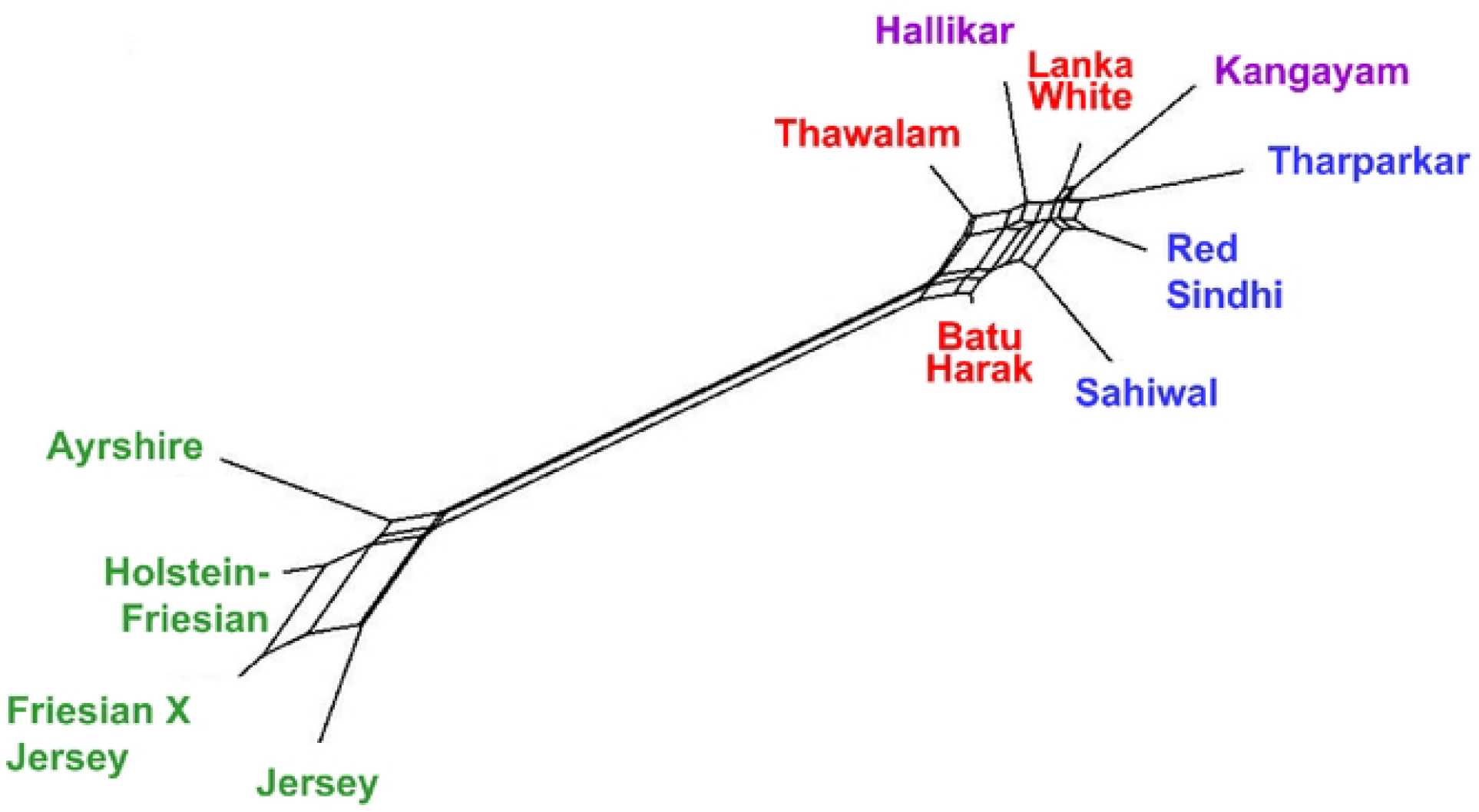
NeighborNet (below) tree derived from pairwise Nei’s genetic distance among Sri Lankan, Indus Valley, South Indian Zebu and com,nercial taurine cattle.

**Figure 3.**
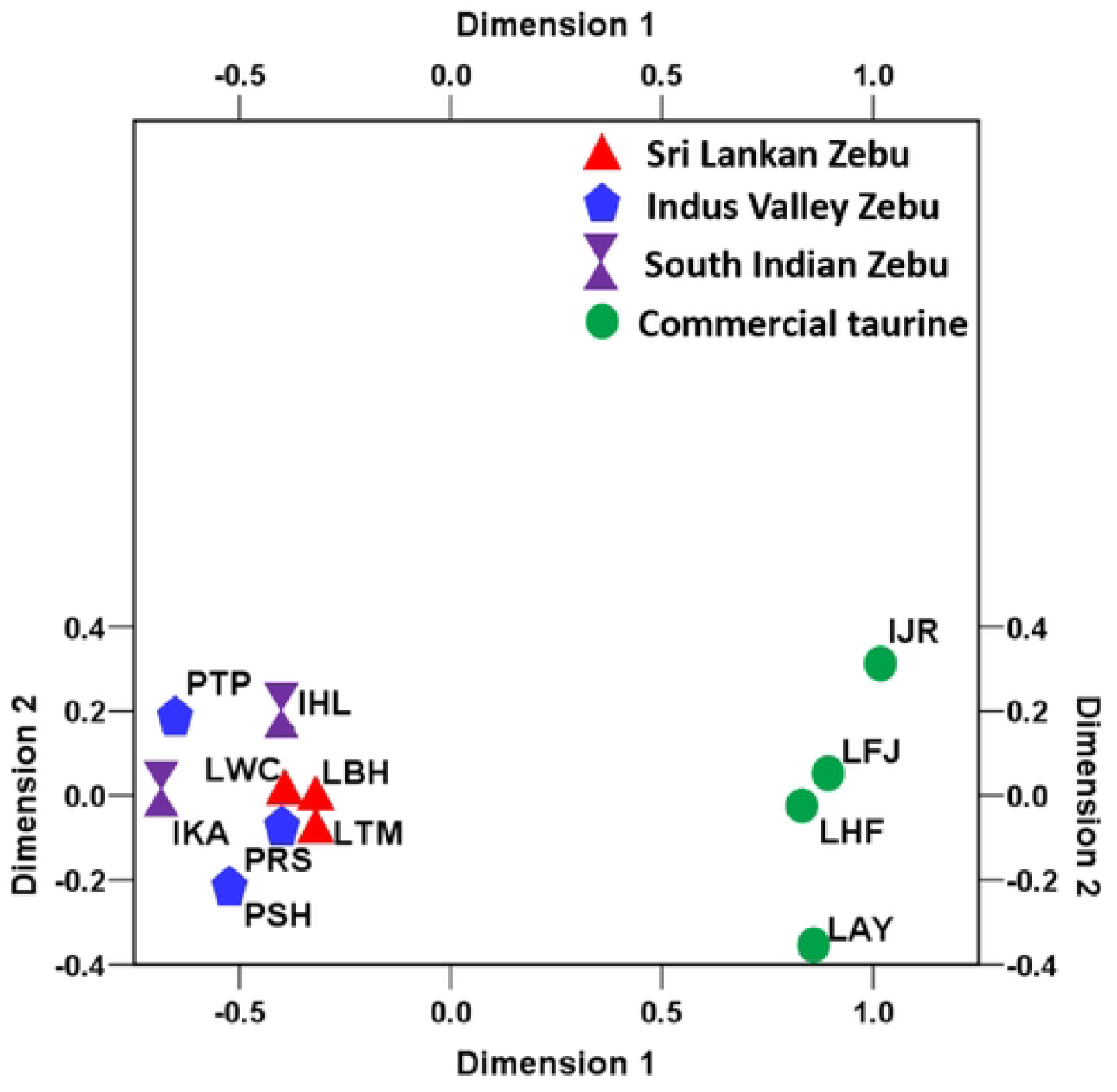
Multi-dimensional scaling display ofpaiiwise F_ST_ among Sri Lankan, South Indian and Indus valley zebu cattle

#### 3.3. Genetic relationships and population structure of Sri Lankan cattle

To further assess the underlying genetic relationship among the investigated zebu cattle populations and to test and validate the results of phylogeny and MDS analysis, analysis of molecular variance (AMOVA) was performed. When no grouping of zebu cattle was assumed, AMOVA revealed 93.93% of total genetic variation was due to differences among individuals within breeds and only 6.07% was accounted for by differences between breeds (Table 3). When zebu breeds were grouped according to geographical origin (Grouping I – Indus valley, South Indian and Sri Lankan zebu), among group variance was 1.0% (P>0.05) while between breed differences within groups accounted for 5.29% (P<0.05). When Sri Lankan zebu were grouped with Indus valley zebu (Grouping II), among group variance was not statistically significant (P>0.05). Similarly, when Sri Lankan zebu was grouped alongside South Indian cattle (Grouping III), among group variance slightly decreased and statistically not significant (P>0.05). When Red Sindhi cattle (Indus valley) alone was grouped with Sri Lankan zebu (as visualized in MDS plot of pairwise FST), among group variance was only 0.86% (P>0.05). However, when Hallikar cattle alone (South India) was grouped with Sri Lankan zebu, among group variance was 2.54% and statistically significant (P<0.01). The results clearly indicated strong genetic relationship of Sri Lankan zebu cattle with Hallikar, an ancestral Mysore type South Indian breed known for its excellent draught characteristics. Bayesian clustering of individuals without prior population information was performed to assess the cryptic genetic structure and degree of admixture among the investigated cattle breeds. Among K = 2 to 14, our dataset was best described with K = 2 genetic clusters, at which the second order rate of change in LnP(D) was maximum (Supplementary Figure SF2). When K = 2 was assumed, the investigated breeds got clustered along taurine-indicine divide (Figure 4). When K = 4 was assumed, Sahiwal cattle clustered separately from other zebu cattle. At K=5, Kangayam cattle clustered separately while at K=6, Hallikar and Tharparkar formed separate clusters (Table 4).

**Figure 4.**
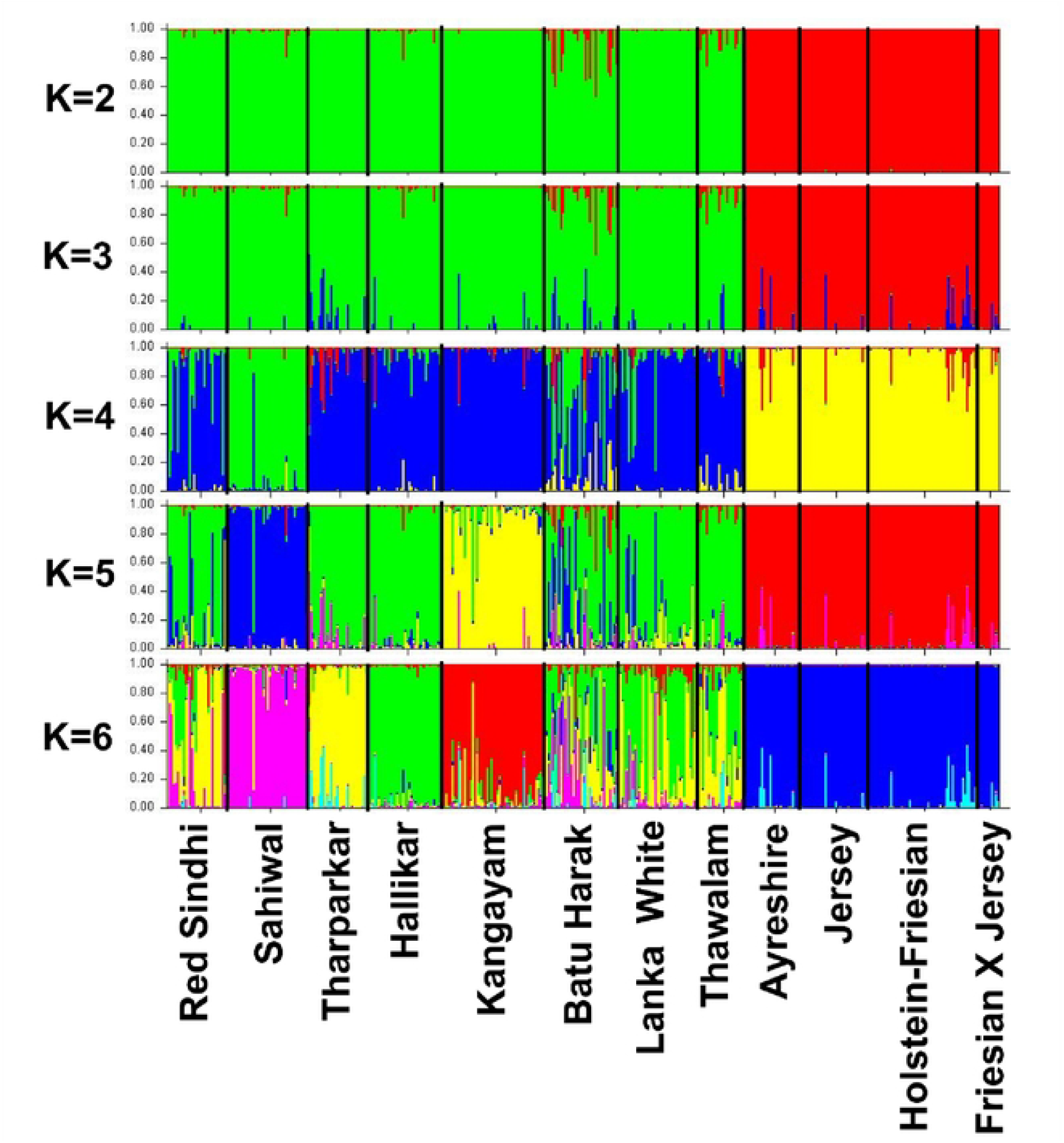
Bayesian clustering of 406 cattle under assumption of 2 to 8 clusters without a priori population information. The population names are given below the box plot with the individuals of different populations separated by vertical black lines

#### 3.4. Mitochondrial DNA diversity in Sri Lankan cattle

The haplogroup classification and diversity of Sri Lankan zebu cattle are presented in Table 5. Two major zebu haplogroups, I1 and I2 were observed in Sri Lankan zebu with the former predominating the later in all the three breeds. A total of 112 haplotypes were observed in the studied breeds, of which 50 haplotypes (34 belonged to I1 lineage, 12 belonged to I2 lineage and two belonged to T3 lineage) were found in Sri Lankan zebu cattle. The haplotype diversity was highest in Batu Harak cattle (0.958) while it was lowest in White cattle (0.802) among the Sri Lankan zebu breeds. The haplotype diversity of I1 lineage was higher than that of I2 lineage in all the three Sri Lankan zebu cattle. Among the haplotypes observed in all the zebu cattle (Sri Lankan, Indus valley and South Indian), 16 were found to be shared across 2 or more breeds, of which 10 belonged to I1 lineage and 6 belonged to I2 lineage (Supplementary Table ST2). A total of 69 singleton haplotypes were observed, of which 44 belonged to I1 lineage and 25 belonged to I2 lineage. Among the I1 lineage, haplotype BWRSPHK_H19_I1 was shared among all the investigated breeds except Thawalam cattle. In case of I2 lineage, haplotype WTRPHK_H106_I2 was shared among all the studied breeds with the exception of Batu Harak and Sahiwal cattle. The diversity of mtDNA control region in the studied cattle populations is presented in Table 6. Overall, 103 polymorphic sites were observed, of which 31 were singleton sites and 70 were parsimony informative sites. The overall nucleotide diversity was 0.009, 0.004 and 0.003 in Batu Harak, Thawalam and White cattle respectively. The nucleotide diversity of I1 lineage in these three breeds was 0.004, 0.002 and 0.001 respectively while the nucleotide diversity of I2 lineage was 0.003, 0.001 and 0.001 respectively. The average number of nucleotide differences was high in Batu Harak cattle (8.22) while it was low in Thawalam (3.26) and White cattle (3.12).

#### 3.5. mtDNA phylogeny and evolutionary relationship

Phylogenetic analysis was performed on all the 112 unique haplotypes along with reference sequences of each lineage, the results of which is presented in Supplementary Figure SF3. The phylogenetic tree conformed to haplogroup classification with the formation of two major indicine clusters I1 and I2. No major sub-clusters were found within each of these two clusters, but three minor clades (one in I1 and two in I2) each with three or more haplotypes were observed. The first minor clade within I2 cluster included unique haplotypes of Batu Harak, Hallikar, Kangayam and Sahiwal cattle while the second minor clade included unique haplotypes of Lanka White and Red Sindhi cattle. The minor clade within I1 cluster included three unique haplotypes of Hallikar and Red Sindhi cattle. The pairwise F_ST_ among zebu cattle from Sri Lanka, Indus valley and South India was estimated utilizing the frequency of mtDNA haplotypes (Table 7). Analysis of I1 haplotypes revealed pairwise F_ST_ varying between zero (Hallikar-Red Sindhi; Sahiwal-Tharparkar) and 0.202 (Thawalam-Lanka White) while I2 haplotypes showed pairwise F_ST_ varying between zero (Hallikar-Sahiwal) and 0.178 (Thawalam-Tharparkar). Among the Sri Lankan zebu breeds, Batu Harak showed low F_STs_ with Tharparkar (F_ST_=0.030) and Red Sindhi (F_ST_=0.039) among I1 haplogroup while it showed lowest F_ST_ with Sahiwal (F_ST_=0.021) among I2 haplogroup. Lanka White cattle showed low F_STs_ with Sahiwal (F_ST_=0.030) and Hallikar (F_ST_=0.050) among I1 haplogroup while it showed low F_STs_ with Red Sindhi (F_ST_=0.020) and Hallikar (F_ST_=0.053) among I2 haplogroup. Interestingly, I1 haplogroup of Thawalam cattle showed high F_STs_ not only with Indus valley and South Indian cattle (F_ST_=0.108-0.199) but also with other Sri Lankan zebu (F_ST_=0.168-0.202) breeds. However, among I2 haplogroup, Thawalam cattle showed low F_STs_ with Sahiwal (F_ST_=0.020) and Red Sindhi (F_ST_=0.039).

#### 3.6. Mismatch distribution, haplotype network and tests for selective neutrality

The demographic history of Sri Lankan zebu cattle was explored by assessing mismatch distribution of nucleotide differences among haplotypes within I1 and I2 maternal lineages (Figure 5). All the three breeds showed unimodal distribution indicating population expansion in both the lineages. The modal peak was observed at three and four nucleotide differences in Batu Harak cattle for I2 and I1 lineages respectively, while the modal peak was observed at two nucleotide differences in Thawalam cattle for both the lineages. In case of Lanka White cattle, haplotype pairs with no differences were abundant followed by pairs with one and two nucleotide differences in both I1 and I2 lineages. The sum of squared differences (SSD) was statistically significant (P<0.01) for I1 lineage of Thawalam and I2 lineage of Batu Harak cattle (Table 8). Although high SSD was observed for I2 lineage of Thawalam cattle, it was not significant statistically (P>0.05). Similarly, raggedness index was significantly high for I1 lineage of Thawalam cattle (P<0.01) and I2 lineage of Batu Harak cattle (P<0.05). The results of neutrality tests revealed negative Tajima’s D values for both the lineages of Batu Harak (P>0.05) and White cattle (P>0.05) indicating an excess of low frequency polymorphisms and demographic expansion. Thawalam cattle showed positive Tajima’s D for both I1 and I2 lineages although statistically not significant (P>0.05). Fu’s FS statistics was negative for both lineages in all the three Sri Lankan native cattle populations and statistically significant (P<0.02). The results indicated the presence of excess number of alleles as would be expected from a recent population expansion or from genetic hitchhiking. Median Joining (MJ) network of control region haplotypes of Sri Lankan, Indus valley and South Indian zebu was reconstructed. The haplotype network results (Figure 6) revealed two distinct lineages, I1 and I2 as expected with expansion events radiating from ancestral nodes. Each of these two ancestral nodes consisted of haplotypes from all the three Sri Lankan native cattle (Batu Harak, Thawalam and White cattle). Interestingly, one of the low frequency nodes formed by haplotypes of Batu Harak and White cattle predated the ancestral and other internal nodes.

**Figure 5.**
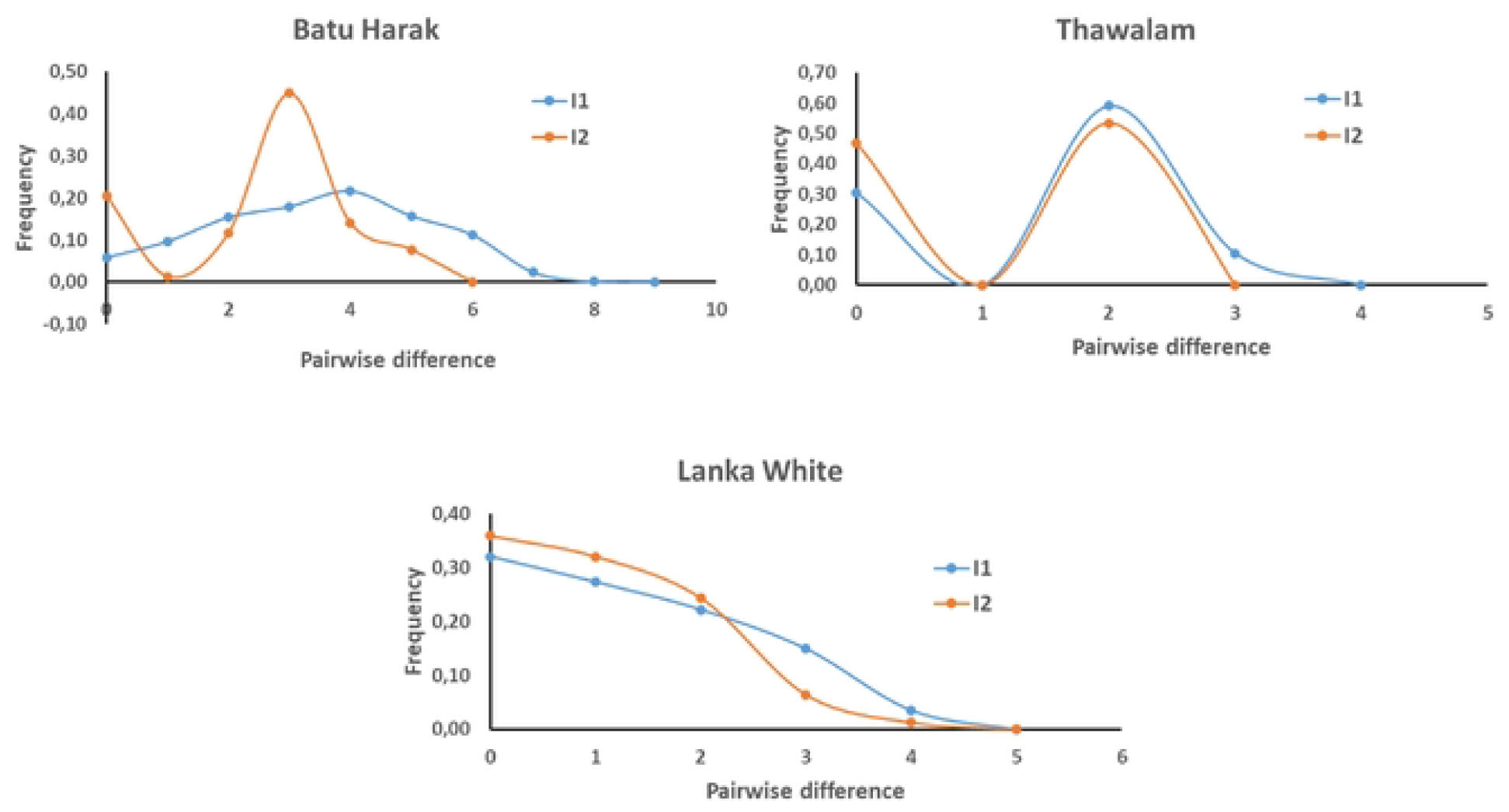
Mismatch distribution of pairwise nucleotide differences among mtDNA control region haplogroups of Sri Lankan zebu cattle

**Figure 6.**
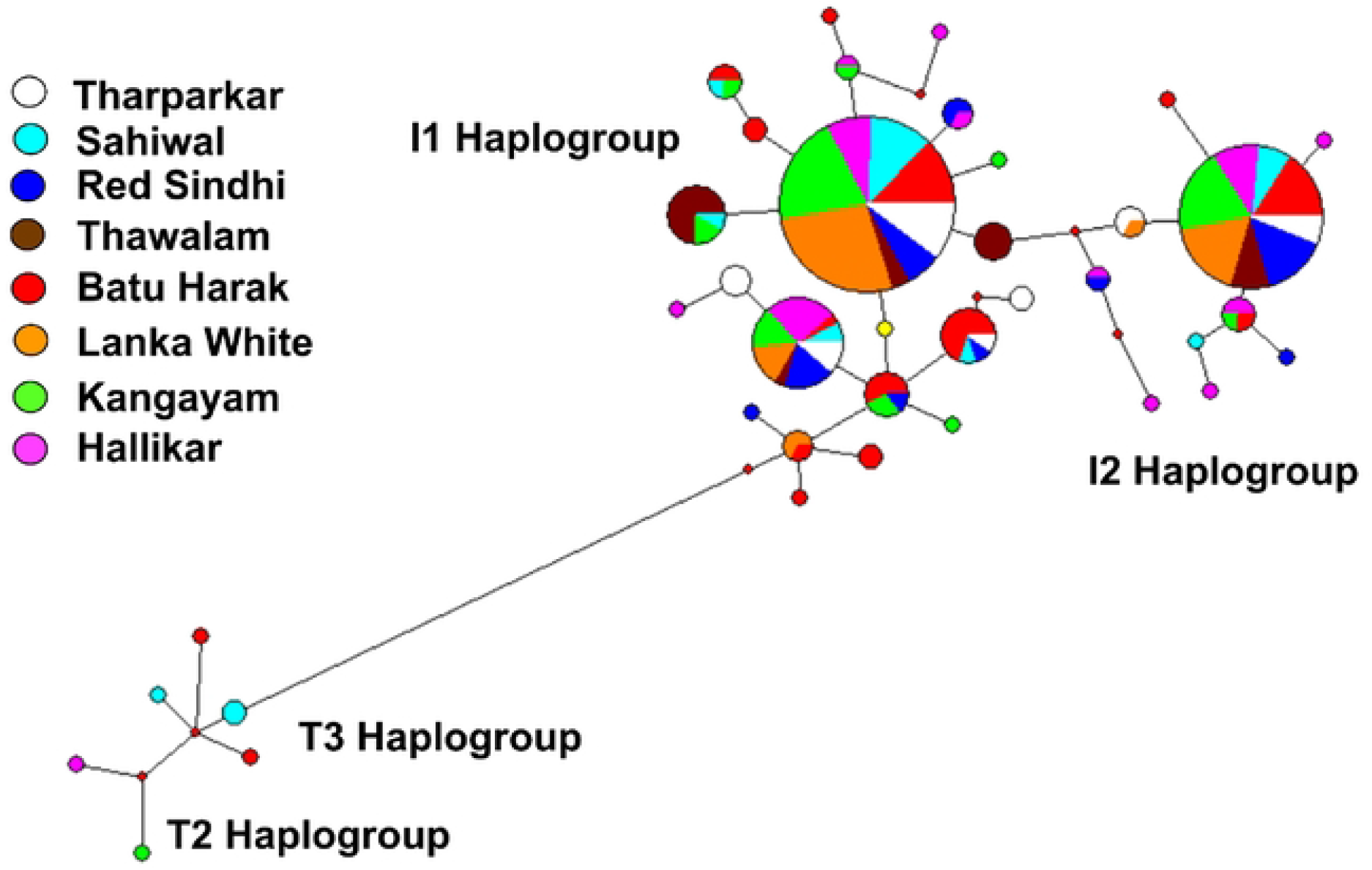
Median Joining network of mtDNA control region haplotypes of Sri Lankan, Indus valley and South Indian zebu (Size of the circle indicates frequency of haplotypes)

## 4. Discussion

### 4.1. Biodiversity and inbreeding estimates

Our study presents the first comprehensive analysis of diversity, genetic and phylogeographic relationship of island cattle of Sri Lanka with zebu cattle of Indus valley and South Indian origin. The existing cattle production system of Sri Lanka comprises of at least five categories of breed types that include native zebu cattle, exotic zebu, native zebu x exotic zebu crosses, exotic taurine, and native zebu x exotic taurine crosses. The distribution of cattle breed types varies with the agro-ecological zones with zebu and zebu crosses dominating the dry zone while purebred exotic taurine and native zebu x exotic taurine crosses dominating the wet zone of the country. The intermediary zone consisted of both zebu x taurine as well as native x exotic zebu crosses (Abeygunawardena et al., 1997). Various breed improvement programs targeting productivity enhancement in indigenous zebu cattle resulted in dilution of germplasm leading to concerns on breed purity and conservation. The present study showed allelic diversity and heterozygosity of Sri Lankan native zebu cattle being slightly higher than the estimates observed for Indus valley (Red Sindhi, Sahiwal and Tharparkar) and South Indian (Hallikar and Kangayam) breeds. Among the Sri Lankan zebu, Batu Harak had the highest n_o,_ H_o_ and H_e_. The extant population of Batu Harak is believed to have resulted from extensive outcrossing of original Lanka cattle with various Indo-Pakistan zebu cattle brought to Sri Lanka (Perera and Jayasuriya 2008; Perera at el., 2010). This might have resulted in higher levels of genetic variability in Batu Harak cattle as compared to other Sri Lankan zebu breeds. The estimates of inbreeding in Sri Lankan native cattle breeds were relatively high and varied from 4.8% (Batu Harak) to 7.2 % (White cattle). Such high estimates of F_IS_ can be associated with heterogeneity of herds sampled within each breed, possibly resulting in Wahlund effect (Hedrick, 2013). Assortative mating can also increase genetic relatedness within the livestock populations and this nonrandom mating pattern can result in increase in heterozygosity deficit. The Sri Lankan White cattle herds mainly consist of animals with similar phenotypes (body size, coat colour, etc.) and farmers preferred bulls with white coat color to ensure the uniformity of herd. The past and current farming practice in the Eastern province (native breed tract of White cattle) also indicates use of few breeding bulls within each herd. Similarly, Thawalam is an isolated population of local zebu in Central and Uva provinces with little gene flow from outside. This might have contributed to consanguineous mating and higher levels of inbreeding within these populations.

### 4.2. Breed history, genetic relationships and population structure

The present-day Lankan cattle (Batu Harak) cattle are believed to represent the descendants of crosses between original archaic “Lanka cattle” and various Indus valley (Indo-Pakistan) zebu breeds (Perera and Jayasuriya 2008; Silva et al., 2008). The White cattle, also called as Thamankaduwa are believed to have originated from the mixture of cattle that existed in ancient Sri Lanka and Indian white cattle breeds (Silva et al., 2008). Anecdotal evidence indicates White cattle probably originated from South India and were brought by tobacco planters in the north-eastern region of the country (Felius, 1995; Nadheer, 2005; Silva et al., 2008). The Thawalam cattle are pack animals that are bred and reared in isolated pockets of Central and Uva provinces of Sri Lanka. Basically, they represent non-descript type, but several generations of breeding in isolation resulted them in emerging as a separate breed with distinct body conformation and capable of pulling loads in hilly regions (Silva et al., 2008). The results of the present study confirmed the close genetic relationships of indigenous Sri Lankan cattle with South Indian and Indus valley zebu, particularly with Hallikar and Red Sindhi cattle respectively. The pairwise genetic differentiation of Sri Lankan zebu cattle from these two breeds were relatively low as compared to other Indus valley and South Indian breeds analyzed in this study. The analysis of molecular variance revealed significant among group variance when the Sri Lankan zebu cattle were grouped with Hallikar cattle (Table 3). All the three Sri Lankan zebu breeds shared significant levels of ancestry with Hallikar cattle breed, while admixture of Indus valley breeds was also noticeable. The proportion of membership coefficient assigned to Hallikar cluster was 39.0%, 50.2% and 37.4% for Batu Harak, Lanka White and Thawalam cattle respectively when K=6 was assumed under unsupervised Bayesian clustering analysis. Similarly, the proportion of membership coefficient assigned to Tharparkar/Red Sindhi cluster was 24.6%, 32.2% and 49.0% while the assignment to Sahiwal cluster was 19.9%, 8.5% and 2.7% in Batu Harak, Lanka White and Thawalam cattle respectively (Table 4). The White cattle showed a relatively high proportion of shared ancestry with South Indian zebu while Thawalam cattle showed high levels of shared ancestry with Indus valley zebu. Batu Harak cattle showed more or less similar levels of shared ancestry with both South Indian and Indus valley zebu cattle. The results of genetic structure analysis revealed strong genetic relationships between Sri Lankan and South Indian zebu cattle, while varied levels of admixture was observed with Indus valley zebu cattle. Introgression of Indus valley zebu in Sri Lankan cattle appears to be a recent phenomenon with the introduction of grading up scheme during 1950s in the mid and low country dry zones using Tharparkar, Red Sindhi and Sahiwal cattle. This was evident from the presence of Sahiwal introgression at specific locations in the native tract of Batu Harak cattle.

### 4.3. Breed purity and taurine introgression

Historically, with the arrival of Europeans, temperate *Bos taurus* breeds were brought into Sri Lanka and since then crossbreeding is used as one of the methods for genetic improvement of milk production in the country. However, indiscriminate breeding of local cattle using semen of taurine and crossbred bulls resulted in dilution of native germplasm. Bayesian clustering analysis was performed to assess the taurine admixture in Sri Lankan zebu cattle using purebred Jersey, Ayrshire and Holstein-Friesian as reference genotypes. The proportion of membership coefficient obtained for individual animals in the inferred zebu and taurine clusters were utilized to estimate taurine admixture and purity of the investigated zebu cattle breeds (Manomohan et al., 2021). The results revealed high levels of breed purity in Lanka White cattle with all the sampled individuals showing >97% zebu ancestry in them. Interestingly, the farmers in the Eastern provincial region predominantly follow natural service for breeding their cattle and the artificial insemination coverage is less than 5% (Abeygunawardena et al., 1997; Sinniah and Pollott, 2006). It is also believed that the White cattle are descendants of the ancient royal herds and white colour is considered as a sign of cleanliness and prosperity (National Livestock Breeding Policy guidelines and strategies for Sri Lanka, 2010). Further, the farmers owning the White cattle herds are aware of the breed purity and use true to the type bulls for breeding, with white coat colour as a primary criterion for selection. With respect to Batu Harak cattle, >38% of animals showed at least 6.25% taurine admixture. Further, more than one-fifth of Batu Hark cattle sampled in the present study showed at least 25% taurine admixture although they had zebu features morphologically. In case of Thawalam cattle, 29.2% of sampled individuals had at least 6.25% taurine admixture while 4.2% had at least 25% taurine admixture. The National Livestock Breeding Policy of Sri Lanka (National Livestock Breeding Policy guidelines and strategies for Sri Lanka, 2010) recommends grading up of native cattle in the low and mid country dry zones using Ayrshire and Jersey breeds with up to 50% of taurine blood level in the crossbreds for improved milk production. The level of taurine admixture in Batu Harak and Thawalam cattle is relatively higher, possibly due to well established AI programs and readily available exotic taurine semen for insemination.

### 4.4. Maternal lineage and phylogeography

The mtDNA maternal lineages of indigenous Sri Lankan zebu cattle belonged to I1 and I2 haplogroups typical to other South Asian zebu cattle (Chen et al., 2010; Manomohan et al., 2021). The mtDNA diversity of Sri Lankan zebu cattle in terms of nucleotide diversity, average number of nucleotide differences and haplotype diversity was relatively low for both I1 and I2 lineages with the exception of Batu Harak cattle. In case of Batu Harak cattle, the diversity parameters were comparable to that of Indus valley, North and South Indian zebu cattle (Chen et al., 2010; Sharma et al., 2015; Manomohan et al., 2021). The mismatch analysis to explore demographic history of Sri Lankan zebu cattle indicated unimodal distribution and potential population expansion in all the three breeds. The SSD and raggedness index was non-significant in both the lineages of all the three breeds except for I1 lineage of Thawalam cattle (P<0.01) and I2 lineage of Batu Harak cattle (P<0.05). The SSD and raggedness index values were relatively high for I2 lineage of Thawalam cattle as well but were not statistically significant (P>0.05). Non-significant values of SSD indicate the goodness of fit between observed and expected data (in a parametric bootstrap simulation analysis) under the population expansion model. Significant values of SSD indicate the deviation of observed data from expected data which is common in populations under demographic equilibrium. Similarly, the values of raggedness index will be larger for populations in demographic equilibrium while they will be small and non-significant for populations under demographic expansion. Interestingly, Tajima’s D statistics was positive for both I1 and I2 lineages of Thawalam cattle, while it was negative in Batu Harak and White cattle. The negative Tajima’s D was significant (P<0.05) only for I2 lineage of White cattle. The Tajima’s D statistics was based on frequency spectrum of mutations while Fu’s FS was based on haplotype distribution. The negative Tajima’s D signifies an excess of low frequency polymorphisms than expected, indicating either expansion of population size or purifying selection. Similarly, negative FS is evidence for an excess number of alleles as would be expected from a recent population expansion or genetic hitchhiking. The results of the present study thus indicated purifying selection among the Sri Lankan zebu cattle breeds. Further, Fu’s FS statistic, sensitive to the presence of singletons and devised more specifically to detect demographic expansion was statistically significant (P<0.01) in all the investigated cattle breeds. However, it is worth to mention that analysis of additional samples from Thawalam cattle will give more insights into its demographic history as it shows signs of population equilibrium.

## Conclusion

The present study revealed moderately high levels of genetic diversity in Sri Lankan zebu cattle breeds. The estimates of inbreeding were relatively high in Thawalam and White cattle breeds. The Sri Lankan zebu cattle breeds showed strong genetic relationships with Hallikar cattle, an ancient breed considered to be ancestor for most Mysore type draught cattle breeds of South India. Recent introgressions of Indus valley zebu like Red Sindhi, Tharparkar and Sahiwal was also evident in Sri Lankan zebu, especially in Batu Harak and Thawalam cattle breeds. mtDNA analysis showed two maternal lineages, I1 and I2 in Sri Lankan zebu cattle with relatively high level of haplotype diversity in Batu Harak cattle. Genetic admixture analysis revealed high levels of breed purity in Lanka White cattle while significant taurine admixture was observed in Batu Harak and Thawalam cattle. Genetic dilution of native zebu germplasm is a cause for concern from biodiversity and conservation standpoint. It is recommended that national breeding organizations consider establishing conservation units for the three native cattle breeds to maintain breed purity and initiate genetic improvement programs. Considering the superior adaptability characteristics of these animals, such an initiative will be beneficial in the long run to develop locally adapted dual-purpose cattle for milk and meat production in Sri Lanka.

## Conflict of interest

The authors report no conflicts of interest.

## Acknowledgment

We like to thank International Atomic Energy Agency, Technical Cooperation project (SRL5046) and University of Peradeniya research grant RG/AF/2013/46/V for financial support and technical assistance. We thank local farmers and veterinarians in Sri Lanka and Pakistan along with Livestock and Dairy Development Departments of different provinces for their help in sampling. The authors also acknowledge the support of S. Bandara, N. Shanjayan, R. Rupasinghe, D.C.A. Gunawardena and staff of Department of Basic Veterinary Sciences, University of Peradneiya, Sri Lanka for their support during the study.

## Notes

### Competing Interest Statement

The authors have declared no competing interest.

